# RUNX proteins desensitize multiple myeloma to lenalidomide via protecting IKZFs from degradation

**DOI:** 10.1101/423350

**Authors:** Nan Zhou, Alvaro Gutierrez Uzquiza, Xiang Yu Zheng, Dan Vogl, Alfred L. Garfall, Luca Bernabei, Anita Saraf, Laurence Florens, Michael P. Washburn, Anuradha Illendula, John H. Bushweller, Luca Busino

## Abstract

Ikaros family zinc finger protein 1 and 3 (IKZF1 and IKZF3) are transcription factors that promote multiple myeloma (MM) proliferation. The immunomodulatory imide drug (IMiD) lenalidomide promotes myeloma cell death via Cereblon (CRBN)-dependent ubiquitylation and proteasome-dependent degradation of IKZF1 and IKZF3. Although IMiDs have been used as first-line drugs for MM, the overall survival of refractory MM patients remains poor and demands the identification of novel agents to potentiate the therapeutic effect of IMiDs. Using an unbiased screen based on mass spectrometry, we identified the Runt-related transcription factor 1 and 3 (RUNX1 and RUNX3) as interactors of IKZF1 and IKZF3. Interaction with RUNX1 and RUNX3 inhibits CRBN-dependent binding, ubiquitylation and degradation of IKZF1 and IKZF3 upon lenalidomide treatment. Inhibition of RUNXs, via genetic ablation or a small molecule (AI-10-104), results in sensitization of myeloma cell lines and primary tumors to lenalidomide. Thus, RUNX inhibition represents a valuable therapeutic opportunity to potentiate IMiDs therapy for the treatment of multiple myeloma.

---

The major therapeutic goal for multiple myeloma (MM) treatment is complete remission and prolonged survival ^1,2^. Small molecules such as the proteasome inhibitor bortezomib and the immunomodulatory imide drugs (IMiDs) thalidomide and lenalidomide, in addition to autologous hematopoietic stem-cell transplantation, have markedly improved overall survival. However, patients with disease refractory to both IMiDs and bortezomib have a median event-free-survival and overall survival of only 5 and 9 months, respectively ^1,2^. Thus, the search for novel agents to potentiate IMiDs represents an important therapeutic goal.

IKZF1 and IKZF3 are transcription factors of the IKZF family, including also IKZF2, 4 and 5 proteins ^3^, that function as homo- and hetero-dimers to regulate lymphopoiesis ^4,5^. Their expression is initiated during the early stages of lymphoid progenitors and increases as lymphocytes differentiate into mature cells ^4^. Genetic data has revealed that the loss of IKZF1 and IKZF3 results in a block of lymphoid lineage differentiation and a susceptibility to develop acute lymphoblastic leukemia (ALL) ^6^. Consistently, IKZF1 and IKZF3 are frequently mutated tumor-suppressor genes in ALL ^7^. In contrast, MM cells display dependency on IKZF1 and IKZF3 for cell-autonomous proliferation ^8,9^. Indeed, the loss of IKZFs by shRNAs or by expression of a dominant-negative IKZF3 mutant inhibits myeloma growth ^8,9^.

The IMiDs thalidomide, lenalidomide and pomalidomide bind to a specific pocket in the E3 ligase Cereblon (CRBN) ^10,11^, promoting CRBN interaction with the IKZF1 and IKZF3 proteins and favoring their ubiquitylation and proteasome-dependent degradation ^8,9^. IMiD-induced degradation of IKZFs promotes the expression of the interferon gene program essential to repress MYC and proliferation of MM cells ^8,9,12,13^. Despite the clinical success of the IMiDs, refractory disease still presents a therapeutic challenge.

In addition to the IKZF family, another family of transcription factors implicated in human hematological cancers is the Runt-related transcription factor family (henceforth referred to as RUNXs). RUNX1 alterations, including translocations, mutations, and gene amplifications, are frequently observed in human leukemia ^14-16^. The RUNX proteins (RUNX1, RUNX2 and RUNX3) are transcription factors that bind to promoters and enhancers via the Runt homology domain (RHD), a domain also required for interaction with its partner CBFβ ^17^. RUNX1 targets multiple genes that are pivotal regulators of hematopoiesis, including the hematopoietic-specific member of the E-twentysix (ETS) family ^18^. Although RUNX1 is essential for murine embryonic development and fetal liver hematopoiesis, conditional deletion of *Runx1* revealed a nonessential role in adult hematopoiesis ^19^.

Here, we show that RUNX1 and RUNX3 physically interact with IKZF1 and IKZF3 *in vitro* and *in vivo*. When complexed with RUNXs, IKZFs become refractory to CRBN-dependent ubiquitylation and degradation induced by IMiDs. Importantly, genetic loss or chemical inhibition of RUNX proteins result in enhanced sensitivity of MM cells to IMiDs. Our data open the possibility of utilizing RUNX inhibition to potentiate IMiD therapy in MM.

## Results

### IKZF1 and IKZF3 physically associate with RUNX1 and RUNX3 in multiple myeloma

IKZF1 and IKZF3 are transcription factors highly expressed in myeloma that contribute to myeloma cell survival ^8,9^. To better understand the function of IKZFs, we sought to identify novel physiologic binding partners. To this end, we generated a human MM cell line, ARP1, stably expressing physiologic levels of FLAG-tagged human IKZF1 or IKZF3 via retroviral delivery. FLAGpeptide eluates from anti-FLAG affinity purifications, either from nuclear extract (nucleoplasm) or benzonase-extracted detergent-insoluble fraction (DNAbound), were trypsinized and subjected to mass spectrometry analysis for analysis of protein identity (Fig. 1a and Supplementary Table 1). Relative to FLAGimmunoprecipitates from ARP-1 cells infected with an empty virus, the peptides corresponding to the NuRD and SWI/SNF complexes, known members of the IKZF1 and IKZF3 complexes, were identified (Fig. 1b, c) ^30,31^. Surprisingly, we identified the transcription factors RUNX1, RUNX3, and CBFβ as interactors of both IKZF1 and IKZF3 (Fig. 1b, c).

**Fig. 1.**
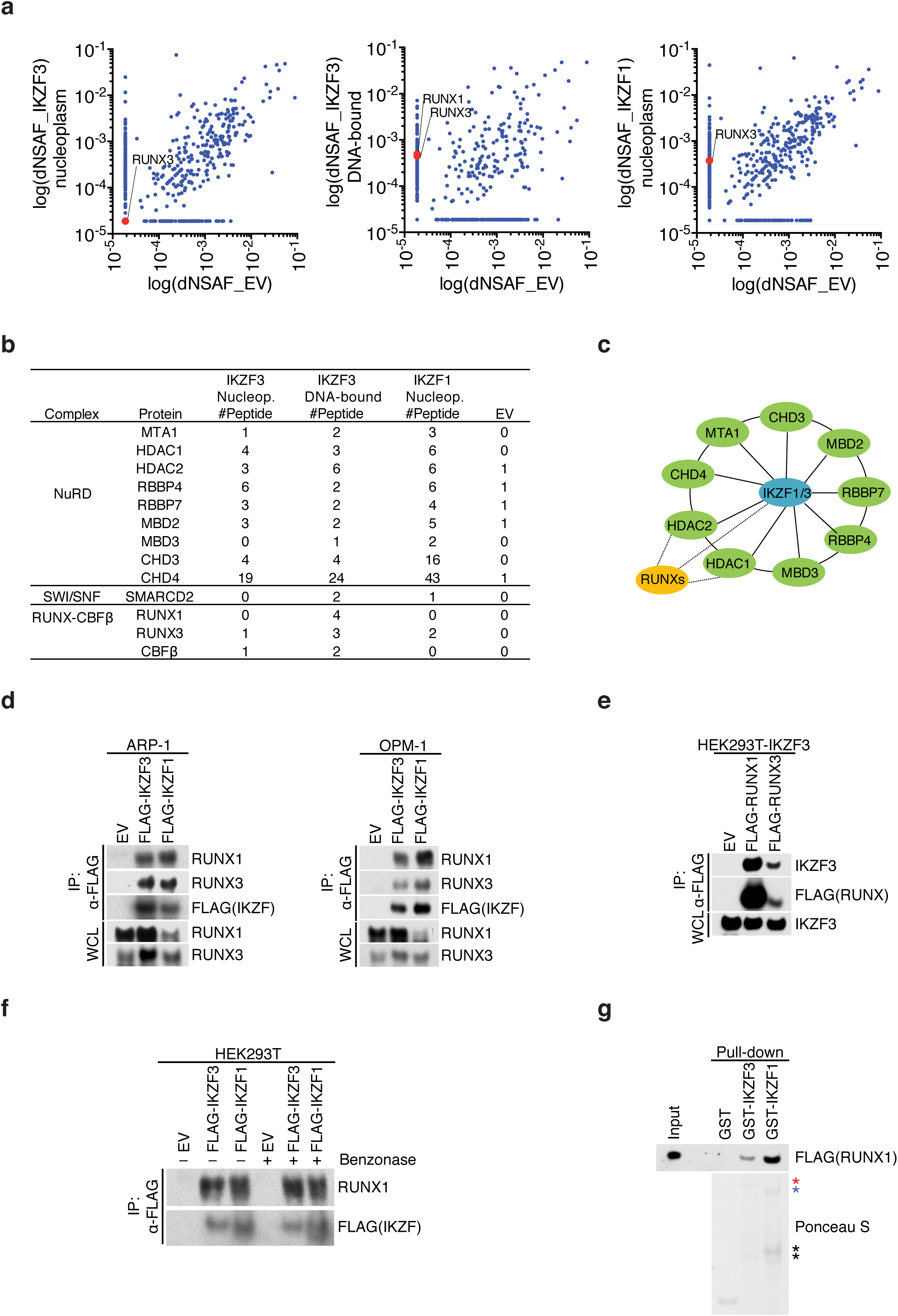
IKZF1 and IKZF3 physically associate with RUNX1 and RUNX3 in multiple myeloma. **(a)** Scatter plot of distributed normalized spectral abundance factor (dNSAF) in FLAG-immunoprecipitates (IP) from ARP-1 cells stably expressing FLAG-tagged IKZF1 and IKZF3 upon mass spectrometry analysis. For proteins with NSAF=0, the lowest NSAF value was arbitrarily assigned. **(b)** List of peptides for the indicated proteins. EV, empty vector. **(c)** Schematic model of IKZF1/3 interactors. For a complete list of interacting proteins see Supplementary Table 1. **(d)** The cell extracts of ARP-1 (left) and OPM-1 (right) cells stably expressing FLAG-tagged IKZF1 and IKZF3 were immunoprecipitated with an anti-FLAG resin and the immunocomplexes were probed with antibodies to the indicated proteins. Specificity of RUNX1 and RUNX3 antibodies was assessed using siRNAs against RUNX1 or RUNX3 (Supplementary Figure 1a). **(e)** HEK293T cells stably expressing IKZF3 were transfected with FLAG-RUNX1 or RUNX3. FLAG-immunoprecipitates were probed with antibodies to the indicated proteins. **(f)** HEK293T cells were transfected with FLAG-tagged IKZF1 or IKZF3. The cell extracts were subjected to anti-FLAG IP and the immunocomplexes were treated with Benzonase for 30 min where indicated. IPs were probed with antibodies to the indicated proteins. **(g)** Purified GST-tagged proteins as indicated were incubated with *in vitro* translated FLAG-tagged RUNX1. GST pull-downs were probed with anti-FLAG antibodies. Ponceau S staining shows the expressions of GST-proteins. The red asterisk indicates GST-IKZF1, blue asterisk shows GST-IKZF3, and black asterisks show cleavage products. Unless otherwise noted, immunoblots are representative of three independent experiments.

To validate our proteomic screen, we performed co-immunoprecipitation experiments in both ARP-1 and OPM-1, two MM cell lines, and confirmed the interaction between stably expressed FLAG-IKZF1 and IKZF3 and endogenous RUNX1 and RUNX3 (Fig. 1d). Reciprocal coimmunoprecipitation of endogenous IKZF3 was also observed in FLAGRUNX1 and RUNX3 immunoprecipitates (Fig. 1e). Antibodies against the endogenous RUNX1/3 and IKZF1/3 were validated by siRNAs (Supplementary Fig. 1a). To rule out the possibility of DNAmediated interaction, we incubated the anti-FLAG-IKZF1 and IKZF3 immunoprecipitates with benzonase to hydrolyze any residual DNA contamination. After extensive washes, we found that RUNX1 was still associated with the IKZFs, suggesting that DNA did not mediate this interaction (Fig. 1f). In agreement with the latter point, purified recombinant IKZF1 and IKZF3 displayed efficient interaction with *in vitro*-translated RUNX1 (Fig. 1g).

Since RUNX1 binds CBFβ ^32,33^, we determined whether IKZFsRUNX1 form a ternary complex with CBFβ (Supplementary Fig. 1b). The interaction between IKZF1 and CBFβ was detected when RUNX1 was co-expressed, suggesting that the three proteins could form a complex. Importantly, the association of CBFβ with IKZF1 was not dependent on the ability of IKZF1 to form homodimers. Indeed, expression of FLAG-IKZF1 (1400), a dimerization-impaired mutant that contains the first four zinc fingers (ZnFs) only ^34^, retained its ability to interact with CBFβ in a RUNX1-dependent manner (Supplementary Fig. 1c).

Collectively, these results suggest that IKZFs assemble a novel complex via direct interaction with RUNXs in MM cells. Moreover, IKZF1 utilizes the DNA-binding ZnFs to assemble a ternary complex with RUNX1-CBFβ.

### IKZF1 and IKZF3 utilize the N-terminal ZnF domain to associate with the activation and inhibition domain in RUNX1

The IKZF proteins contain four N-terminal C2H2-type ZnFs involved in sequence-specific DNA binding ^30,34-36^, and two additional Cterminal ZnFs that promote homo- and hetero-dimerization with the IKZF family members ^34,37^. To determine the domain required for interaction with RUNX1, we generated a series of N-terminal and C-terminal deletion mutants in IKZF3 (Fig. 2a, b). While deletion of the C-terminal ZnFs involved in dimerization did not result in loss of RUNX1 interaction (mutants Δ420-509 and Δ480-509) (Fig. 2a, b), deletion of the N-terminal ZnFs eliminated IKZF3-RUNX1 association (mutants Δ1-181, Δ1-243 and Δ1-279) (Fig. 2a, b).

**Fig. 2.**
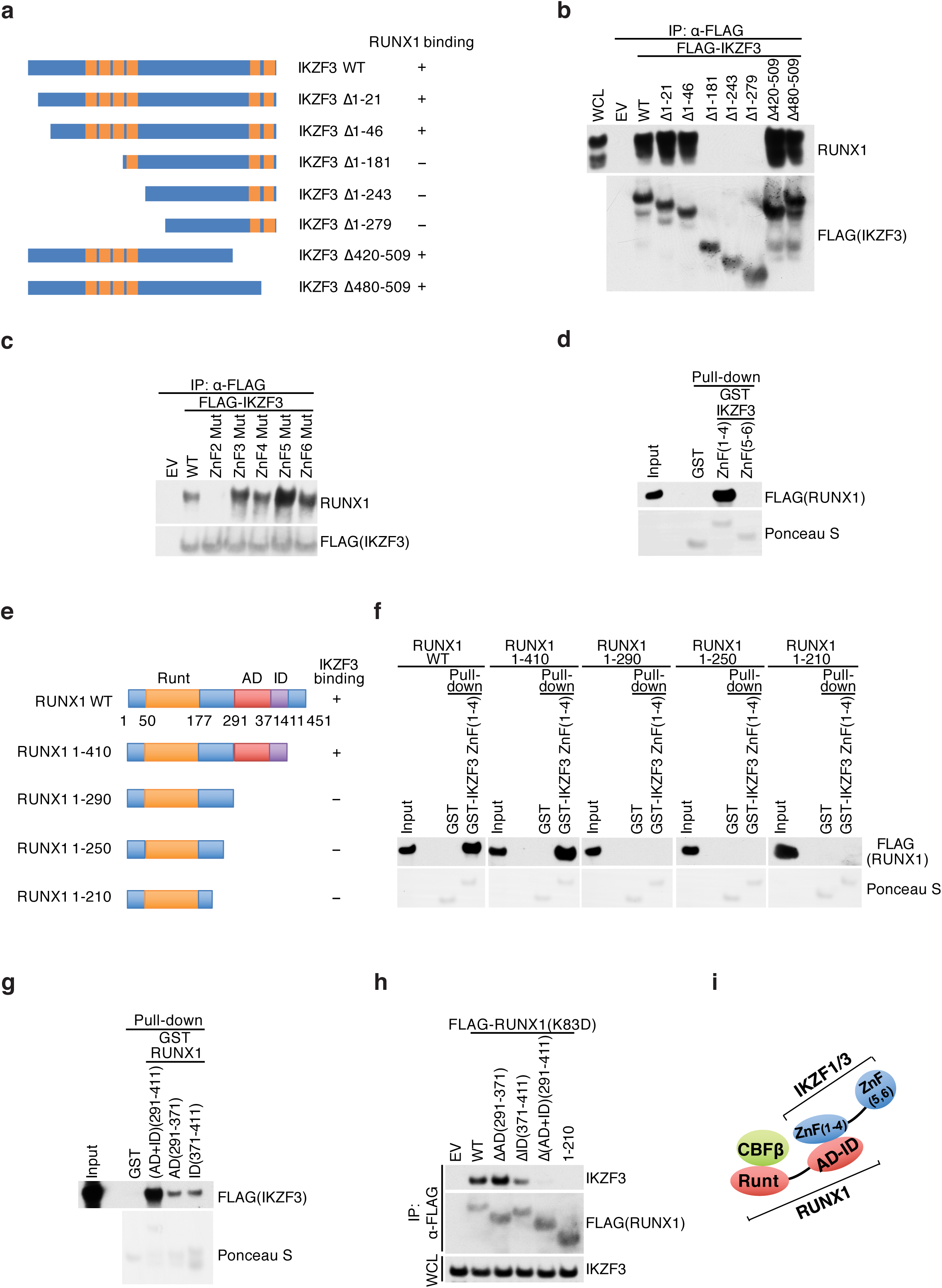
IKZF1 and IKZF3 utilize the N-terminal ZnF domain to associate with the activation and inhibition domain of RUNX1. **(a)** HEK293T cells were transfected with constructs encoding an empty vector (EV), FLAG-tagged IKZF3 wild-type (WT), or mutants. A schematic representation of IKZF3 mutants is shown. IKZF3 mutants that interact (+) or do not interact (-) with RUNX1 are shown. **(b)** Immunoblot analysis of FLAG-IKZF3 immunoprecipitation (IP). Immunocomplexes were probed with antibodies to the indicated proteins. **(c)** HEK293T cells were transfected with constructs encoding an empty vector (EV), FLAG-tagged IKZF3 wild-type (WT) or mutants as indicated. The cell extracts were subjected to anti-FLAG IP and the immunocomplexes were probed with antibodies to the indicated proteins. **(d)** Purified GST-tagged proteins as indicated were incubated with *in vitro* translated FLAG-tagged RUNX1. GST pull-downs were probed with anti-FLAG antibodies. Ponceau S staining shows the expressions of GST-proteins. **(e)** Schematic representation of RUNX1 mutants. RUNX1 mutants that interact (+) or do not interact (-) with IKZF3 are shown. **(f)** Immunoblot analysis of GST pull-downs. Purified GST-tagged proteins as indicated were incubated with *in vitro* translated FLAG-tagged RUNX1. GST pull-downs were probed with anti-FLAG antibodies. Ponceau S staining shows the expressions of GST-proteins. **(g)** Same as in *(f)*, except that the indicated GST-proteins were used. **(h)** HEK293T cells stably expressing IKZF3 were transfected with constructs encoding an empty vector (EV), FLAG-tagged RUNX1 wild-type (WT) or mutants as indicated. The whole cell lysates (WCL) were subjected to anti-FLAG IP and the immunocomplexes were probed with antibodies to the indicated proteins. **(i)** Schematic model of the interaction between IKZF1/3 and the AD and ID of RUNX1. Unless otherwise noted, immunoblots are representative of three independent experiments.

In an effort to understand the relevance of individual ZnFs, we assessed the interaction of IKZF3 with mutations in the zinc-coordinating cysteines of each ZnF to RUNX1 (Fig. 2c). With this analysis, we demonstrated that ZnF (2) is critical in mediating interaction with RUNX1. To further confirm that IKZF3 N-terminal ZnFs are directly involved in RUNX1 interaction, we purified IKZF3 containing ZnF (1-4) or ZnF (5-6) and tested the association with RUNX1 *in vitro*. In agreement with our *in vivo* mapping (Fig. 2a), IKZF3-ZnF (1-4) are sufficient to directly interact with RUNX1, in contrast to IKZF3-ZnF (5-6), which are dispensable (Fig. 2d). Notably, the ZnF (1-4) in IKZF3 are highly similar to the corresponding ZnFs in IKZF1, with ZnF (2) displaying 100% sequence homology (Supplementary Fig. 1d). Thus we interpret that binding results would be identical amongst IKZF1 and IKZF3. Attempts to recapitulate binding *in vitro* utilizing recombinant single ZnFs resulted in minimal interaction with RUNX1 (data not shown), potentially suggesting that the interaction surface with RUNX1 may have multiple contact points beyond ZnF (2).

RUNX1 contains an N-terminal conserved DNA-binding domain (called the Runt domain). This region is responsible for both sequence-specific DNA-binding and hetero-dimerization with the core binding factor β subunit (CBFβ)^32,33^. At the C-terminus, RUNX1 possesses an activation domain (AD) that interacts with transcriptional co-activators such as p300/CBP ^38^ and an inhibitory domain (ID) that counteracts the activation domain ^39^. In an attempt to determine the IKZF3-interacting domain of RUNX1, we performed *in vitro* Glutathione S-Transferase (GST)-pull down utilizing the IKZF3 ZnF (1-4) fragment fused to GST as bait against *in vitro*-translated RUNX1 (either full-length or Cterminal truncations) (Fig. 2e, f). Deletion of a C-terminal region containing the AD and ID resulted in ablation of interaction with IKZF3, suggesting that AD and/or ID are necessary to promote interaction with IKZF3. To assess sufficiency, we purified recombinant AD and ID (either individually or in combination) fused to GST (Fig. 2g). Both GST-AD and GST-ID were sufficient to interact with IKZF3 *in vitro*, although when the AD and ID were combined, the interaction with IKZF3 was greatly enhanced (Fig. 2g). To confirm the requirement for the AD and ID in cells, we performed co-immunoprecipitation experiments using lysates of HEK293T cells expressing constructs encoding FLAG-tagged RUNX1 (Fig. 2h). To minimize non-specific interactions mediated by DNA, we expressed RUNX1 (K83D), a mutant which is incapable of binding DNA ^40^. In agreement with our *in vitro* mapping, simultaneous ablation of AD and ID resulted in the loss of IKZF3 interaction (Fig. 2e, f), while individual deletion of the AD or ID did not impair, or only partially impaired, the interaction with IKZF3, respectively. Both *in vitro* and *in vivo* mapping data suggest that the interaction surface on RUNX1 requires the cooperation of the AD and ID.

Together, our data suggest that IKZF1 and IKZF3 utilize the DNA-binding N-terminal ZnFs but not the C-terminal ZnFs involved in homo- and hetero-dimerization, to interact with the AD and ID of RUNX1 (Fig. 2i).

### RUNXs inhibit CRBN-dependent IKZF1 and IKZF3 degradation

Earlier work has suggested that the therapeutic effects of the IMiDs reflect a CRBN gain-of-function favoring degradation of IKZFs ^8,9^. The region of IKZF1 that mediates lenalidomide-dependent binding to CRBN is located within the ZnF (2) ^8,9^, thus overlapping with the identified RUNX1 binding site. These findings led us to hypothesize that RUNXs compete with CRBN for binding to IKZFs.

To test this hypothesis, we first analyzed the levels of IKZF1 in OPM-1 cells depleted of the RUNX proteins. In OPM-1 *RUNX1*^*+/+*^ *RUNX3*^*+/+*^ cells, lenalidomide induced degradation of IKZF1 but not RUNX1 and RUNX3 (Fig. 3a). While the kinetics was unchanged in the single *RUNX1*^*-/-*^ or *RUNX3*^*-/-*^ knock-out cells, the double knock-out cells displayed an overall accelerated degradation of IKZF1 (Fig. 3a). Importantly, IKZF1 and IKZF3 were the only CRBN-substrates that displayed enhanced degradation in the *RUNX1*^*-/-*^ *RUNX3*^*-/-*^ cells upon drug treatment; ZFP91, another CRBN-target^41^, showed a slight time-dependent decrease when cells were treated with lenalidomide and no change in the *RUNX1*^*-/-*^ *RUNX3*^*-/-*^ cells (Fig. 3b).

**Fig. 3.**
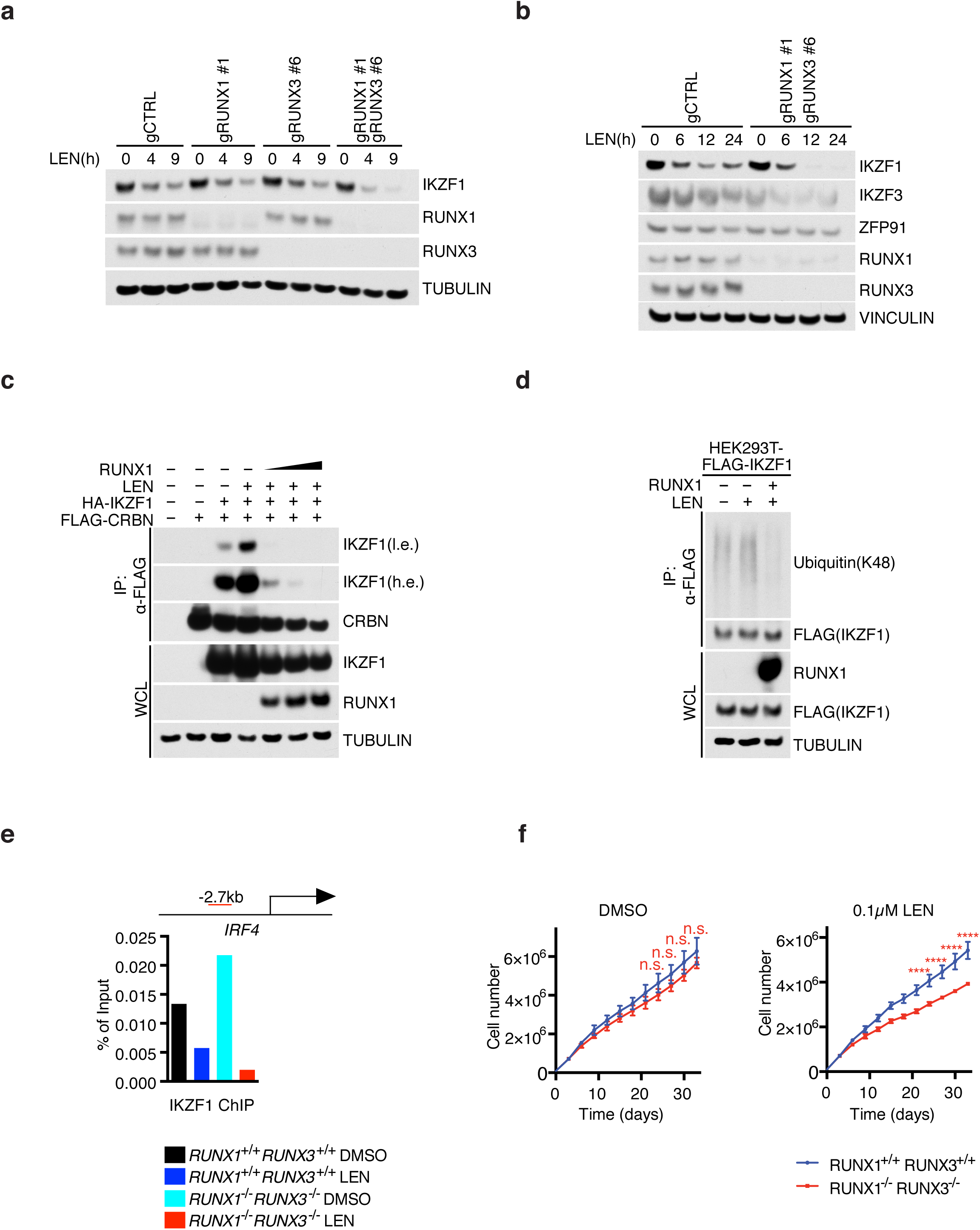
RUNXs inhibit CRBN-dependent IKZF1 and IKZF3 degradation. **(a)** Immunoblot analysis of whole cell lysates from OPM-1 cells treated with 1μM lenalidomide for the indicated durations of time. **(b)** Same as in *(a)* except that OPM-1 cells were treated with lenalidomide for the indicated time points. **(c)** HEK293T cells were transfected with FLAG-CRBN, HA-IKZF1 and un-tagged RUNX1. Where indicated, cells were treated with 2 μM lenalidomide for 6 hours before harvesting. The cell extracts were subjected to anti-FLAG IP and the immunocomplexes were probed with antibodies to the indicated proteins. A low exposure (l.e.) and high exposure (h.e.) are shown for IKZF1. **(d)** Same as in *(c)* except that HEK293T cells were transfected with FLAG-IKZF1 and un-tagged RUNX1. **(e)** Chromatin immunoprecipitations of IKZF1 in OPM-1 cells coupled with qRT-PCR using primers for *IRF4* promoter under the indicated conditions. Red bar shows the distance between the primers and transcription start site of *IRF4*. (n=2 independent experiments). **(f)** Cell counts of GFP/Cherry-sorted OPM-1 *RUNX1*^*+/+*^ *RUNX3*^*+/+*^ or *RUNX1*^*-/-*^*RUNX3*^*-/-*^ cells grown in media containing DMSO or 0.1 μM lenalidomide (mean±s.d., n=3 independent experiments, two-way ANOVA, n.s., not significant, \*\*\*\**P*≤0.0001). Unless otherwise noted, immunoblots are representative of three independent experiments.

To gain mechanistic insights into how RUNX proteins protect IKZF1 from IMiD-dependent degradation, we set out to test whether RUNX1 competes out CRBN for binding to IKZF1. In HEK293T cells treated with lenalidomide, IKZF1 was effectively enriched in FLAG-CRBN immunoprecipitates (Fig. 3c). Importantly, overexpression of RUNX1 abrogated IKZF1 binding to CRBN, in line with the hypothesis that RUNX1 and CRBN compete for the same binding site in IKZF1. RUNX1 was able to not only displace IKZF1-CRBN interaction, but also block CRBN-dependent IKZF1 ubiquitylation (Fig. 3d).

Previous evidences revealed that lenalidomide down-regulates *IRF4* transcription, connecting this mechanism to the antimyeloma activity ^42-45^. Thus, we assessed whether the dimer RUNX1-IKZF1 is changed by lenalidomide at the IRF4 locus by chromatin immunoprecipitation (ChIP) (Fig. 3e). We confirmed decreased binding of IKZF1 at the *IRF4* locus by ChIP in OPM-1 cells treated with lenalidomide. In *RUNX1*^*-/-*^ *RUNX3*^*-/-*^ cells, although slightly increased in DMSO treated cells, IKZF1 downregulation by lenalidomide was more profound, confirming the function of RUNXs in protecting IKZF1 from degradation. This data suggests that RUNXs and IKZFs work together at similar genomic sites and that ablation of RUNX proteins results into an amplification of the lenalidomide effect towards IKZF1/3 inhibition.

Next, we assessed the sensitivity of MM cell lines to lenalidomide. Since the clinical efficacy of IMiDs is associated with CRBN expression in MM ^46^, we first assessed protein levels of CRBN in three MM cells lines and found that ARP-1 expressed low CRBN as opposed to OPM-1 and KMS-11 cells (Supplementary Fig. 2a). Accordingly, OPM-1 and KMS-11 cells displayed lenalidomide-sensitivity when grown in media replenished with fresh drug every three days, while ARP-1 cells were insensitive (Supplementary Fig. 2b). Importantly, while proliferation of OPM-1 *RUNX1*^*-/-*^ *RUNX3*^-/-^ cells was not changed as compared to *RUNX1*^*+/+*^ *RUNX3*^*+/+*^ cells in absence of drug, a significant decrease in proliferation was detectable in the knockout cells upon treatment with lenalidomide (Fig. 3f). Notably, these cells displayed sensitivity to 0.1 µM lenalidomide, a concentration at which a minimal effect on MM proliferation was detectable (Supplementary Fig. 2b).

Altogether, these data suggest that RUNX proteins protect IKZFs from CRBN-dependent degradation. In myeloma, genetic ablation of RUNXs sensitizes cells to low doses of lenalidomide.

### RUNX inhibition dissociates the IKZFs-RUNXs complex and potentiates the cytotoxic effect of lenalidomide in myeloma

To assess whether chemical inhibition of RUNXs results in changes in RUNXs-IKZFs interaction, we tested the pan-RUNX inhibitor, AI-10-104 ^47^. AI-10-104 is designed to interfere with the association of RUNX with CBFβ, thereby leaving RUNX in an auto-inhibited state. AI-10-104 induced a dose-dependent dissociation of CBFβ and IKZF3 from RUNX1 (Fig. 4a) and RUNX3 (Fig. 4b). Importantly, treatment of cells with increasing concentrations of AI-10-104 resulted in increasing RUNX1 and RUNX3 dissociation from IKZF3 (Fig. 4a, b). Similar data were produced when IKZF1 was pulled down (Fig. 4c, d). Thus, the RUNXs-IKZFs complex can be chemically-dissociated with the small molecule, AI-10-104.

**Fig. 4.**
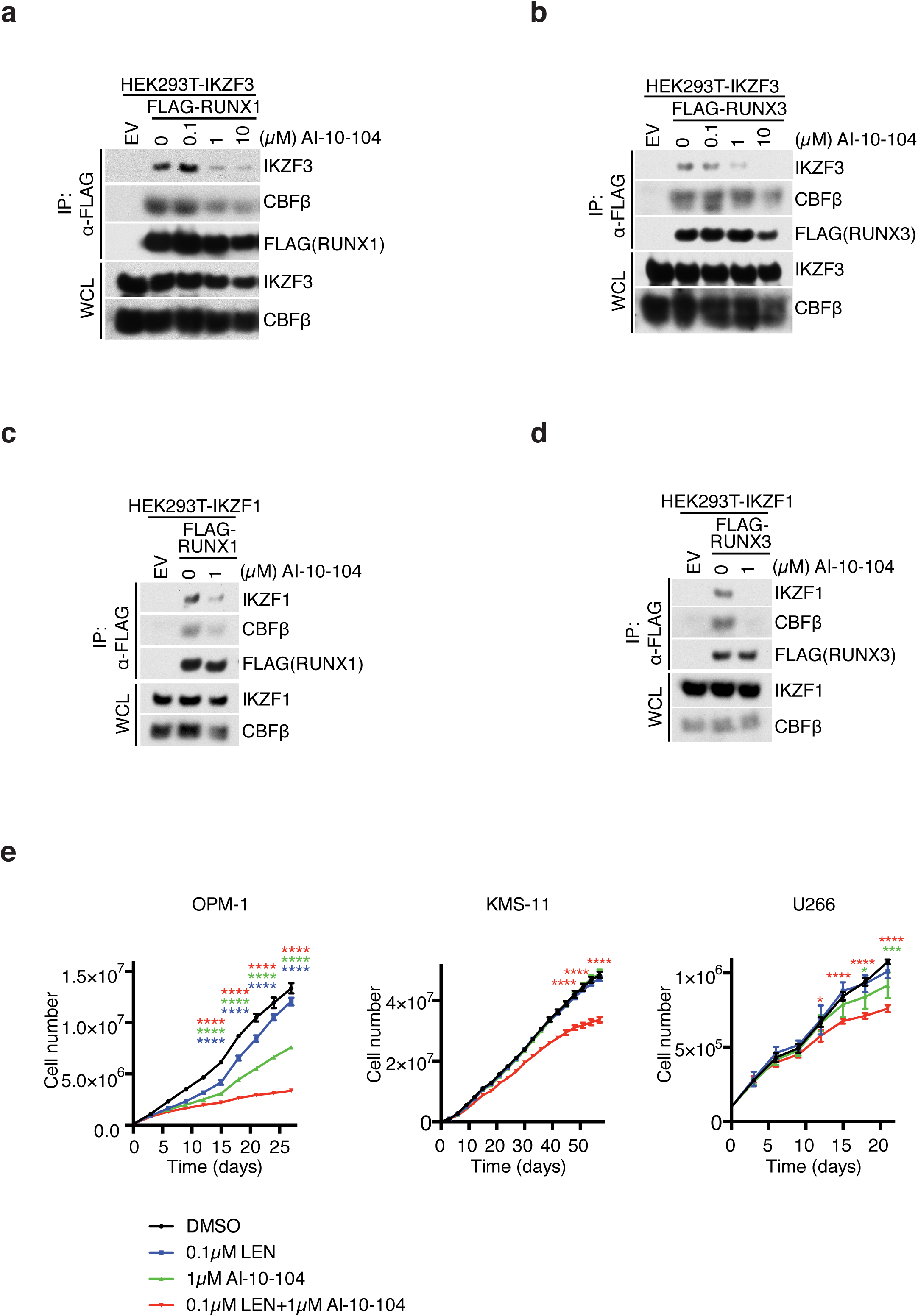
Small molecule AI-10-104 can dissociate the IKZFs-RUNXs complex and potentiate the anti-proliferative effect of lenalidomide. **(a)** HEK293T cells stably expressing IKZF3 were transfected with constructs encoding an empty vector or FLAG-tagged RUNX1 and treated with increasing amounts of RUNX inhibitor (AI-10-104). Immunoprecipitated FLAG-tagged RUNX1 was probed with antibodies to the indicated proteins. **(b)** Same as in *(a)*, except that FLAG-tagged RUNX3 was used. **(c)** HEK293T cells stably expressing IKZF1 were transfected with constructs encoding an empty vector or FLAG-tagged RUNX1 and treated with DMSO or 1 μM RUNX inhibitor (AI-10-104). Immunoprecipitated FLAG-tagged RUNX1 was probed with antibodies to the indicated proteins. **(d)** Same as in *(c)*, except that FLAG-tagged RUNX3 was used. **(e)** Cell counts of the indicated multiple myeloma cell lines grown in media containing the indicated concentration of lenalidomide (LEN), RUNX inhibitor (AI-10-104), and combination (mean±s.d., n=3 independent experiments, two-way ANOVA, \**P* value≤0.05, \*\*\**P* value≤0.001, \*\*\*\**P* value≤0.0001). Unless otherwise noted, immunoblots are representative of three independent experiments.

Next, we tested whether the inhibition of RUNXs could enhance the cytotoxic effect of lenalidomide. To this end, we treated three MM cell lines with low doses of lenalidomide. As previously shown (Supplementary Fig. 2b), low doses of lenalidomide have a modest effect on MM cell proliferation (Fig. 4e). Similarly, treatment of cells with RUNX inhibitor AI-10-104 resulted in moderate (OPM-1 cells) or no (KMS-11 and U266 cells) effect on proliferation when utilized at 0.1 or 1 µM (Fig. 4e), in line with the effect of combined RUNX1/RUNX3 loss on OPM-1 proliferation (Fig. 3f). Importantly, combination of the two drugs induced an overall proliferation defect in the MM cell lines tested (Fig. 4e), suggesting that inhibition of RUNXs proteins potentiates the anti-proliferative effect of lenalidomide, consistent with the finding that lenalidomide promotes enhanced degradation of IKZFs and cytotoxicity in *RUNX1*^*-/-*^ *RUNX3*^*-/-*^ cells (Fig. 3a-e).

Altogether, genetic loss or chemical inhibition of RUNX proteins increases the cytotoxic effect of lenalidomide in MM cell lines.

### RUNX inhibition potentiates the transcriptional response induced by lenalidomide

To understand the molecular basis for the efficacy of the combination regiment of lenalidomide and RUNX inhibition in treating myeloma, we profiled gene expression changes by RNA sequencing of OPM-1 cells (Supplementary Tables 2, 3 and 4). To this end, we treated cells with 0.1 µM lenalidomide or 1 µM AI-10-104 and both in combination. As shown previously, single treatments at the indicated concentrations are not effective in blocking cell proliferation (Supplementary Fig. 2, 3). Notably, low doses of lenalidomide resulted in deregulation of 69 genes as compared to DMSO counterparts while RUNX inhibition resulted in changes in 38 genes. Importantly, the combination of drugs promoted deregulation of 105 genes (Fig. 5a, b and Supplementary Fig. 4) with upregulation of 60 genes specifically (Fig. 5b). Gene ontology (GO) analysis revealed that the most consistent signatures upregulated by combining lenalidomide and RUNX inhibitor were those associated with interferon signaling, immune response, and inflammatory response (Fig. 5c). Similarly, gene set enrichment analysis (GSEA) revealed significant expression of interferon response gene signatures (Fig. 5d). Specifically, OPM-1 cells treated with the combination of drugs upregulated genes related to interferon (IFN) response relative to either agent alone, including *HLA-DQA1, HLA-DRB1, CD74, OAS2, ISG15, NMI, IFIH1*, and *EPSTI1* (Fig. 5e).

**Fig. 5.**
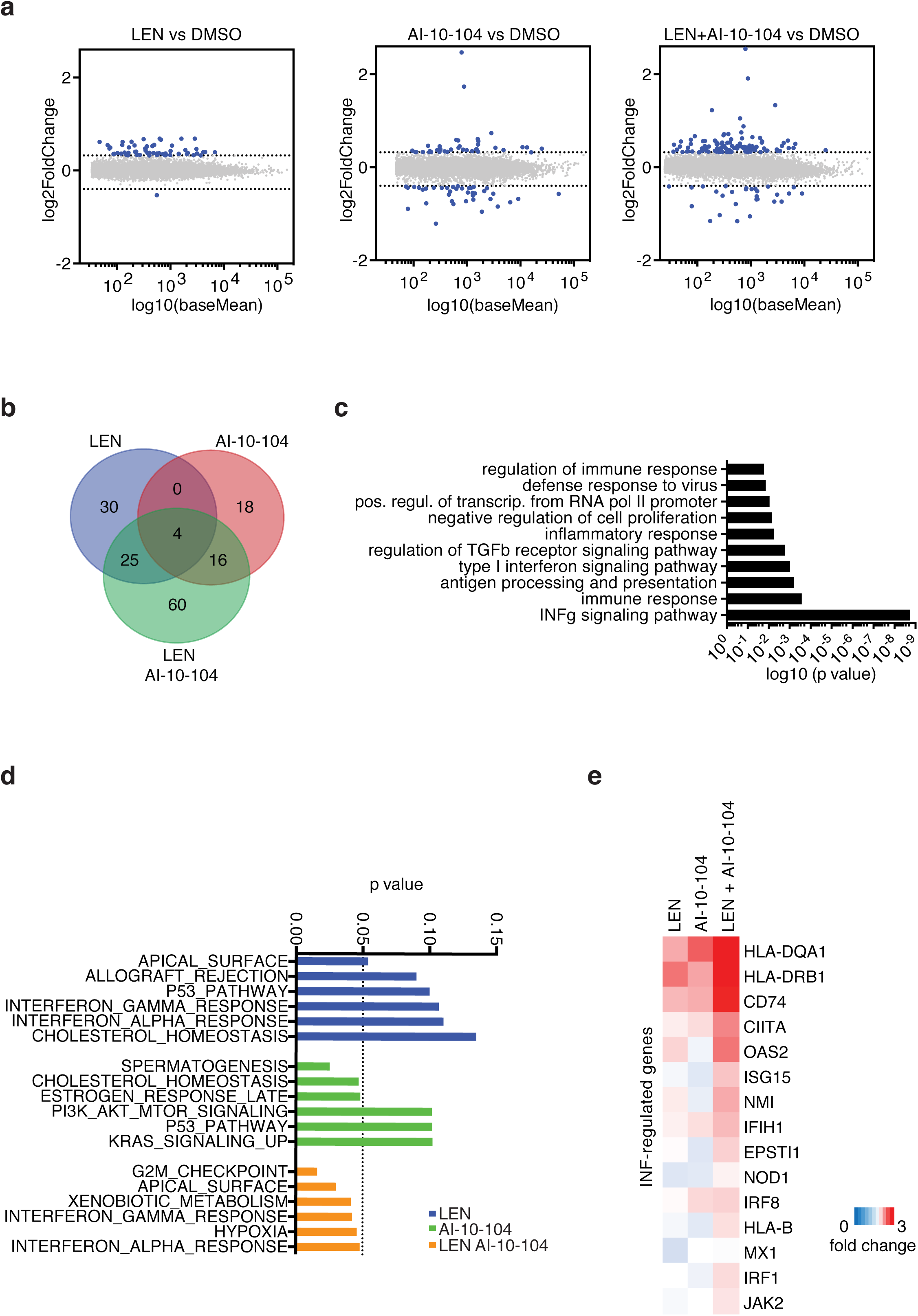
RUNX inhibition potentiates the transcriptional response induced by lenalidomide. **(a)** Fold-change plot of gene expression levels in OPM-1 cells treated with 0.1 μM lenalidomide (LEN), 1 μM RUNX inhibitor (AI-10-104), or combination for 48 hours and compared to DMSO (n=3 independent experiments, DeSeq2). **(b)** Venn diagram showing the overlap of genes up-regulated or down-regulated by lenalidomide, AI-10-104, and combination and compared to DMSO (n=3 independent experiments, *P*≤0.05). **(c)** Gene ontology (GO) analysis of genes regulated by the combination of lenalidomide and AI-10-104. Bar plot for the −log_10_ of the *P* value of the top 10 enriched GO terms of genes regulated is shown. **(d)** *P*-value graph of GSEA enrichment signatures of differentially-expressed genes in OPM-1 cells treated with 0.1 μM lenalidomide (LEN), 1 μM RUNX inhibitor (AI-10-104), or combination and compared to DMSO. **(c)** Heat map showing the relative expression of selected interferon signaling genes in OPM-1 cells treated with 0.1 μM lenalidomide (LEN), 1 μM RUNX inhibitor (AI-10-104), or combination and compared to DMSO (mean, n=3 independent experiments, *P*≤0.05).

These observations, coupled with the fact that myeloma cells are more sensitive to the combination of lenalidomide and RUNX inhibitor treatment prompted us to conclude that inhibition of RUNXs enhances the cellular response to lenalidomide, such that lower doses of lenalidomide are able to gain a therapeutic effect.

### Chemical inhibition of RUNXs potentiates lenalidomide toxicity in primary multiple myeloma cells

We evaluated primary myeloma samples for their sensitivity to combinatorial therapy consisting of RUNX inhibitor and lenalidomide. Primary myeloma cells were isolated from the iliac crest of patients and subjected to CD138-positive purification. Treatment of diagnostic and relapsed myeloma samples (Fig. 6b) with AI-10-104 or lenalidomide alone resulted in minimal or no change in cell viability when compared to DMSO (Fig. 6a). The combinatorial treatment of AI-10-104 and lenalidomide instead displayed a significant inhibitory effect on the viability of primary myeloma samples. These data are in line with our previous observations in MM cell lines, where single treatment at low doses of lenalidomide or AI-10-104 did not reduce cell viability as compared with DMSO (Fig. 4c). Again, combination of the two drugs effectively reduced cell viability, suggesting an additive effect. Of note, the cytotoxic effect of the lenalidomide and AI-10-104 combination regiment was not dependent on patient treatment history (Fig. 6b).

**Fig. 6.**
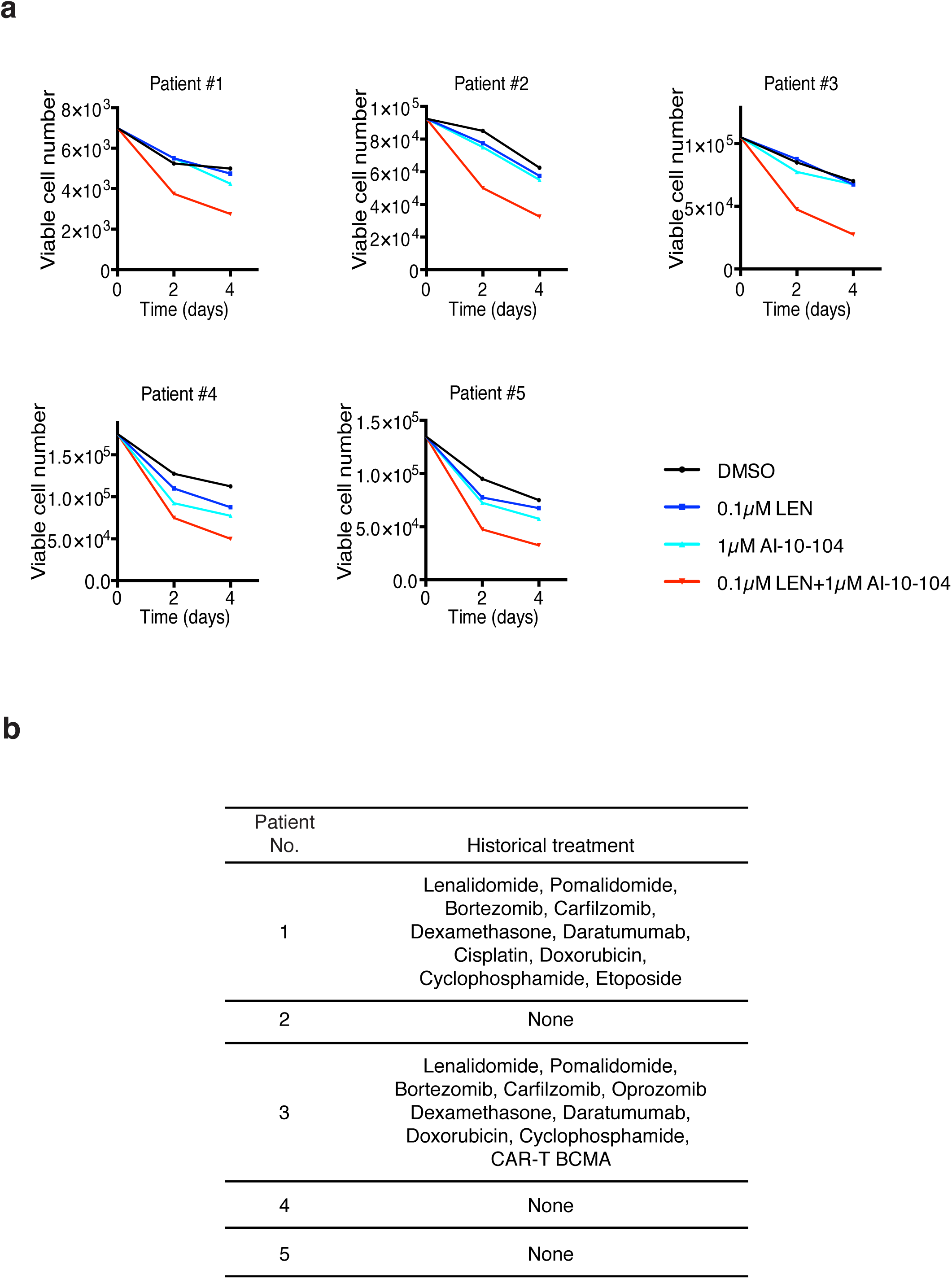
Chemical inhibition of RUNXs potentiates lenalidomide toxicity in primary multiple myeloma cells. **(a)** Primary MM cells were treated with DMSO, 0.1 μM lenalidomide, 1 μM AI-10-104, and combination. Every 2 days, viable cells were counted using Trypan blue exclusion assay **(b)** The historical treatments of five MM patients are shown.

Importantly, treatment of normal human hematopoietic cells with AI-10-104 resulted in an average IC_50_ of ∼15 µM ^47,48^, which greatly exceeded the toxic concentration for MM cell lines and MM primary cells. Thus, RUNX inhibition in myeloma patients should not result in toxic adverse effects on normal hematopoietic stem and progenitor cells. Our data suggest that a therapeutic window may exist for the combinatorial therapy of AI-10-104 derivatives and lenalidomide in MM patients.

## Discussion

Here, we report for the first time that the transcription factors IKZF1 and IKZF3 are physically associated with the RUNX1 and RUNX3 proteins, master regulators of Hematopoiesis ^^^^49^^^^. IKZFs utilize the N-terminal DNA-binding ZnFs to interact with the AD and ID in RUNX1 and form a trimeric complex with RUNX1 binding partner CBFβ. Our findings are particularly interesting when compared with the anti-myeloma effect of the IMiD drug, lenalidomide. Lenalidomide binds CRBN, a CUL4 E3 ubiquitin ligase, and promotes the ubiquitylation and proteolysis of IKZF1 and IKZF3 ^8,9^. We show that interaction with RUNX1 results in the inhibition of IKZF1 ubiquitylation and degradation via the CRBN/lenalidomide complex. Mapping analysis reveals that RUNX1 interacts with the N-terminal ZnFs of IKZF1, a domain also involved in binding to CRBN. Therefore, it is reasonable to conclude that RUNX1 and CRBN compete for IKZF1 interaction and the ablation of RUNX1 and RUNX3 results in enhanced degradation of IKZF1 and IKZF3 upon exposure to lenalidomide. Consequently, *RUNX1*^*-/-*^*RUNX3*^*-/-*^ MM cells display toxicity to nanomolar concentrations of lenalidomide. Importantly, the enhanced degradation of the IKZF proteins in the *RUNX1*^*-/-*^ *RUNX3*^-/-^ cells occurs within 4 to 12 hours; time points at which no difference of cell viability is noticeable.

The cellular signaling programs responsible for CRBN-IMiD-mediated myeloma cytotoxicity are associated with upregulation of the interferon signaling pathway ^8,9,12,42^. In line with the cytotoxic effect in MM cells, RUNX inhibitor potentiates the transcriptional response induced by lenalidomide culminating in the activation of interferon response genes. Thus, deregulation of RUNXs axis could represent an avenue to enhance the anti-myeloma effects of the IMiDs. Further investigation would be necessary to uncover non-cell autonomous mechanisms of this combination therapy; for instance, how lenalidomide and AI-10-104 affect the interaction between myeloma and stroma cells in the bone marrow ^50^.

Introduction of proteasome inhibitors (bortezomib and carfilzomib) and IMiDs have changed the treatment paradigm for myeloma ^51-53^. However, myeloma remains incurable and new treatments are currently being studied. Here, we show that inhibition of RUNXs via the small molecule inhibitor AI-10-104 ^47^ potentiates the cytotoxic effect of lenalidomide in MM cells. RUNX1 downregulation was previously suggested as a possible therapeutic avenue for the treatment of MM ^54^. Furthermore, previous evidence points out a role for RUNX2 in promoting expression of multiple metastatic genes and favoring homing of MM cells to the bone ^55^. Our data suggest that RUNX proteins are dispensable with regard to the proliferation potential of MM cells, and the drugs that achieve pan-inhibition of the RUNX proteins might act as an intervention point for the treatment of MM, particularly in combination with low doses of lenalidomide (Supplementary Fig. 5).

## Methods

### Cell culture

ARP-1, OPM-1, KMS-11, and U266 were maintained in RPMI1640 media containing 10% fetal bovine serum (FBS). HEK293T cells were maintained in Dulbecco’s modified Eagle’s media (DMEM) containing 10% FBS. Primary myeloma cells were obtained from the iliac crest aspirates from MM patients. All patients provided written informed consent under a research sample collection protocol approved by the University of Pennsylvania Institutional Review Board. CD138^+^ cells were isolated with Ficoll Paque Plus (GE Healthcare, #17-1440-02) and CD138^+^ Microbeads (Miltonic Biotic, #130-051-301) according to the manufacturer’s protocol. Primary CD138^+^ cells were maintained in RPMI1640 media containing 50% own serum from the corresponding patients.

### Reagents

The following antibodies were used: anti-FLAG (Sigma, F7425), anti-IKZF3 (Cell Signaling, #15103), anti-IKZF1 (Cell Signaling, #9034), anti-RUNX1 (Abcam, ab92336), anti-RUNX3 (Cell Signaling, #9647), anti-CBFβ (Santa Cruz, sc-20693), anti-CRBN (Sigma, HPA045910), anti-ubiquitin (K48) (EMD Millipore, #05-1307), anti-TUBULIN (Santa Cruz, sc-8035), anti-VINCULIN (Santa Cruz, sc-73614), anti-Rabbit IgG-HRP (GE Healthcare, NA934V), and anti-mouse IgG-HRP (GE Healthcare, NA931V). The following agarose beads were used: anti-FLAG M2 Affinity Gel (Sigma, A2220) and Glutathione Sepharose 4B (GE Healthcare, #17075601). The following *in vitro* translation kit was used: TNT T7 Coupled Reticulocyte Lysate Systems (Promega, L4610). Benzonase was used according to manufacturer (Sigma, E1014). The following compounds were used: Lenalidomide (Sigma, CDS022536), and AI-10-104 was kindly provided by Dr. John H. Bushweller.

### Gene silencing by siRNA

siRNA sequences are as follows: siRUNX1, 5’-CCUCGAAGACAUCGGCAGAAA-3’; siRUNX2, 5’-CUCUGCACCAAGUCCUUUU-3’;and siRUNX3, 5’-CCUUCAAGGUGGUGGCAUU-3’. The following siRNAs were obtained from Dharmacon: siIKZF1 #1 (D-019092-01), siIKZF1 #2 (D-019092-02), siIKZF1 #3, (D-019092-03), siIKZF1 #4 (D-019092-04), siIKZF3 #3 (D-006945-03), siIKZF3 #4 (D-006945-04), siIKZF3 #17 (D-006945-17), siIKZF3 #18 (D-006945-18), and ON-TARGETplus Non-targeting Control siRNAs were obtained from Dharmacon (D-001810-01-20).

### Plasmids

The detailed information regarding plasmid constructions is available by request. And the target sequences used to knock-out human RUNXs are as follows: hRUNX1_gRNA#1 Fwd: ATGAGCGAGGCGTTGCCGCT, Rev: AGCGGCAACGCCTCGCTCAT; hRUNX3_gRNA#6 Fwd: GCCCGAGGTGCGCTCGATGG, Rev: CCATCGAGCGCACCTCGGGC; and control_gRNA Fwd: CTTCGAAATGTCCGTTCGGT, Rev: ACCGAACGGACATTTCGAAG

### Immunoprecipitation and immunoblot

Cells were lysed in NP-40 buffer (0.1% NP-40, 15 mM Tris-HCl pH7.4, 1 mM EDTA, 150 mM NaCl, 1 mM MgCl_2_, 10% Glycerol) containing protease inhibitors (Sigma, #11697498001) and the lysates were incubated with anti-FLAG Gel at 4 °C overnight. After washed, the anti-FLAG Gel was mixed with Laemmli buffer and boiled at 95 °C for 5 min. After SDS-PAGE electrophoresis and transfer, primary antibodies and HRP-linked secondary antibodies were incubated with the membrane for 1 hour at room temperature and overnight at 4 °C, respectively. Where indicated, anti-FLAG immunoprecipitates were resuspended in 50 μl of NP-40 buffer, incubated with 1 μl of Benzonase for 30 min on ice, and further washed twice. For GST pull-down, GST-tagged proteins were incubated with *in vitro* translated FLAG-tagged proteins at 4 °C overnight. After washed with PBS-T three times and PBS once, the membrane was detected by the chemiluminescence system (Thermo Fisher Scientific, #32106).

### Chromatin immunoprecipitation (ChIP)

Cells were cross-linked using 1% formaldehyde for 5 min. Then cells were incubated with 125 mM glycine for 5 min. After centrifugation at 1000 rpm, 4 °C for 5 min, cell pellets were washed with PBS once and lysed with cell lysis buffer (0.2% NP-40, 100 mM Tris-HCl pH8, 10 mM NaCl) for 10 min on ice. After centrifugation at 2500 rpm, 4 °C for 5 min, the pellets were re-suspended with nucleus lysis buffer (0.1% SDS, 50 mM Tris-HCl pH8, 10 mM EDTA) to do sonication using the Covaris S220 system (Thermo Fisher Scientific, #4465653) according to the manufacturer’s protocol. After sonication, the cell lysates were centrifuged at 13,000 rpm, 4 °C for 5 min and the supernatant was collected. Then 10 μl of Dynabeads Protein A (Life Technology, #10001D) were blocked with 1 mg/ml of BSA at 4 °C for 1 hour and 30 μl of Dynabeads were washed with PBS for three times and incubated with 5 μg of antibodies at 4 °C for 4 hours. Three hundred μl of sonicated nucleus lysates were incubated with 10 μl of BSA-blocked Dynabeads at 4 °C for 3 hours. After 3 hours, the pre-cleared 300 μl of nucleus lysates and 700 μl of IP buffer (0.01% SDS, 1% TritonX-100, 20 mM Tris-HCl pH8, 2 mM EDTA, 150 mM NaCl) was incubated with 30 μl of antibody-conjugated Dynabeads at 4 °C overnight. Then the Dynabeads were washed twice with low salt IP wash buffer (0.01% SDS, 1% TritonX-100, 20 mM Tris-HCl pH8, 2 mM EDTA, 50 mM NaCl), twice with medium salt IP wash buffer (0.01% SDS, 1% TritonX-100, 20 mM Tris-HCl pH8, 2 mM EDTA, 300 mM NaCl), and twice with high salt IP wash buffer (0.01% SDS, 1% TritonX-100, 20 mM Tris-HCl pH8, 2 mM EDTA, 500 mM NaCl). The Dynabeads were washed once with LiCl buffer (1% NP-40, 1% deoxycholic acid, 10 mM Tris-HCl pH8, 1 mM EDTA, 0.25 M LiCl) and once with TE buffer (10 mM Tris-HCl pH8, 1 mM EDTA). The chromatin reverse cross-linking and DNA elution was then conducted using IPure kit (diagenode, C03010015) according to the manufacturer’s protocol.

### Quantitative Real-Time PCR (qRT-qPCR)

qRT-PCR was conducted with SYBR Green PCR Master Mix (Thermo Fisher Scientific, #4309155) and the comparative C_T_ method was utilized for relative quantification on the ViiA 7 Real-Time PCR system (Thermo Fisher Scientific). The primer sequences are as follows: *hIRF4* Fwd: 5’-AGTTGCAGGTTGACCTACGG-3’, Rev: 5’-AGCTTTCACCCGTTGAGCTT-3’.

### MudPIT analysis

TCA-precipitates were urea-denatured, reduced, alkylated and digested with endoproteinase Lys-C (Roche), followed by modified trypsin (Roche) ^20, 21^. Peptide mixtures were loaded onto 100-µm fused silica microcapillary columns packed with 5-μm C18 reverse phase (Aqua, Phenomenex), strong cation exchange particles (Luna, Phenomenex)^22^. Loaded microcapillary columns were placed in-line with a Quaternary Agilent 1100 series HPLC pump and a LTQ linear ion trap mass spectrometer equipped with a nano-LC electrospray ionization source (Thermo Scientific). Fully-automated 10-step MudPIT runs were carried out on the electrosprayed peptides, as described ^23.^ Tandem mass (MS/MS) spectra were interpreted using SEQUEST ^24^ against a database of 72,956 sequences, consisting of 72,968 non-redundant human proteins (downloaded from NCBI on 2015-03-25), 148 usual contaminants (such as human keratins, IgGs, and proteolytic enzymes), and, to estimate false-discovery rates, 73,091 randomized amino-acid sequences derived from each non-redundant protein entry. Peptide/spectrum matches were sorted and selected using DTASelect with the following criteria set: spectra/peptide matches were only retained if they had a DeltCN of at least 0.08 and a minimum xcorr of 1.8 for singly, 2.5 for doubly, and 3.5 for triply-charged spectra. In addition, peptides had to be fully tryptic and at least seven amino acids long. Combining all runs, proteins had to be detected by at least two such peptides, or one peptide with two independent spectra. Under these criteria the averagel FDRs at the protein and spectral levels were 0.5%±0.3 and 0.06% ± 0.05, respectively. Peptide hits from multiple runs were compared using CONTRAST ^25^. To estimate relative protein levels, normalized spectral abundance factors (NSAFs) were calculated for each detected protein, as described ^26,27,28^. All mass spectrometry data files may be accessed from the MassIVE repository at ftp://MSV000081712@massive.ucsd.edu.

### Transfection and retrovirus-mediated gene transfer

HEK293T cells were transfected with plasmids using polyethylenimine (PEI) (Polysciences, #24765). For retrovirus and lentivirus production, GP2-293 packaging cells (Clontech) or pCMV-DeltaR8.2 were used. After seventy-two hours of transfection, the virus-containing media was collected and used to spininfect the cells at 1,800 rpm for 40 min. Cells were then incubated with the viral supernatant overnight. For siRNA transfection, the Neon transfection system (Thermo Fisher Scientific, #MPK5000) was used.

### RNA-Seq

Total RNA was extracted from OPM-1 cells using RNeasy Mini Kit (QIAGEN, #74104) and polyA+ transcripts were isolated with NEBNext Poly(A) mRNA Magnetic Isolation Module (NEB, #7490). RNA-Seq libraries were prepared with NEBNext Ultra Directional RNA Library Prep Kit for Illumina (NEB, E7420). Three biological replicates were sequenced on a NextSeq 500 (Illumina) at a depth of at least 2×10^7^ reads each. Reads were mapped and analyzed with a bioinformatic pipeline based on STAR, SAMTOOLS, and the R packages DEGseq and DEseq2. We used human genome version GRCh38. GO analyses were performed using version 6.8 of the DAVID web server. GSEA analyses were performed using pre-ranked GSEA using a weighted scoring^29^.

### Cell proliferation assay

OPM-1, KMS-11, and U266 cells were cultured in 6-well plates. The initial cell number was 100,000 cells per well and the cell numbers were counted and re-plated at the initial concentration every 3 days. Lenalidomide and AI-10-104 were added into the culture media after dilution every 3 days. The cell counting was conducted with Z2 Coulter Counter Analyzer (Beckman Coulter, #6605700).

### Trypan blue exclusion assay

Fifteen μl of 0.4% Trypan blue solution (Cellgro, #25-900-CI) was mixed with 15 μl of CD138^+^ cell suspension and incubated for 2 min at room temperature. The viable cell numbers were counted using a hemocytometer (Fisher Scientific, #02-671-10).

### Statistical analyses

All statistical analyses were performed with Prism 6 (GraphPad), and sample sizes and reproducibility for each figure are shown in the figure legends. Student’s *t*-tests and one-way or two-way ANOVA analyses were performed and indicated in the figure legends. All graphs show mean values with error bars signifying s.d. as indicated in the figure legends.

### Data availability

The authors declare that the data supporting the findings of this study are available upon reasonable requests.

## Acknowledgements

The authors thank Ashley N. Hughes and Grant Grothusen for critically reading the manuscript and Nancy A. Speck for kindly providing RUNX1 constructs. This work was supported in part by grant R00-CA16618104 and R01-CA207513-01 from the National Cancer Institute to L.B.

## Author Contributions

L.B. conceived, directed the project and oversaw the results. N.Z. designed and performed most experiments. X.Z. helped N.Z. with experiments in Fig. 3i, 4c, S2a, S3. A.G.U. performed experiments in Fig. 1a, 1b, 2b, 2c. A.S., L.F. and M.P.W. performed the mass spectrometry analysis of the IKZF complexes purified by A.G.U.. D.V., A.L.G., L.Be. isolated the primary myeloma sample. A.I. and J.H.B. provided with the AI-10-104. L.B. and N.Z. wrote the manuscript.

